# Connectome architecture favours within-module diffusion and between-module routing

**DOI:** 10.1101/2025.02.10.637586

**Authors:** Caio Seguin, Maria Grazia Puxeddu, Joshua Faskowitz, Richard F. Betzel, Olaf Sporns

## Abstract

Connectomes are the structural scaffold for signalling within nervous systems. While many network models have been proposed to describe connectome communication, current approaches assume that every pair of neural elements communicates according to the same principle. Connectomes, however, are heterogeneous networks, comprising elements with varied topological and neurobiological makeups. In this paper, we investigate how connectome architecture may facilitate different signalling regimes depending on the topological embedding of communicating neural elements. Specifically, we test the hypothesis that the modular structure of brain networks fosters a dual mode of communication balancing diffusion—passive signal broadcasting—and routing—selective transmission via efficient paths. To this end, we introduce the relative diffusion score (RDS), a measure to quantify the proportional capacity for network communication via diffusion versus routing. We examined the interplay between RDS and connectome architecture in 6 organisms spanning a wide range of spatial resolutions and connectivity mapping techniques—from the complete nervous system of the larval fly to the inter-areal human connectome. Our analyses establish multiple lines of evidence suggesting that connectomes may be universally organised to support within-module diffusion and between-module routing. Using a series of rewiring null models, we untangle the contributions of connectome topology and geometry to the relationship between routing, diffusion and modular architecture. In conclusion, our work puts forth a hybrid conceptualisation of neural communication, in which diffusion contributes to functional segregation by concentrating information within localised clusters, while specialised signal routes enable fast, long-range and cross-system functional integration.

---

The connectome describes the structural connectivity between elements of a nervous system [1–4]. At every spatial scale, from synapses connecting neurons to white matter fibres intersecting grey matter regions, the connectome is the anatomical substrate for communication between neural elements. Understanding how information is transmitted via the complex wiring of the connectome to support brain function is one of the key challenges of modern neuroscience [5, 6].

A range of computational models seek to capture how the connectome shapes neural communication, including biophysical [7–9], control [10, 11] and graph-theoretical [12, 13] approaches. Here we focus on the later domain, i.e., network communication models that describe connectome signalling using concepts from graph theory and network science [14]. This class of models proposes policies to guide signal propagation in brain networks and quantify the structural capacity for communication between neural elements. Accumulating evidence supports the utility of these models in explaining a wide range of phenomena, including, to name a few, the spread pathogens of over the course of human neurodegenerative illness [15, 16], the propagation of electrical stimulation through the human cortex [17], the functional impact of targeted lesions in the macaque brain [18], and the polysynaptic integration of sensory-motor information in *Drosophila melanogaster* [19, 20].

Network communication models can be placed on a spectrum defined by two opposing endpoints: (i) diffusion processes, characterised by stochastic flow and broadcasting along multiple network fronts, and (ii) routing protocols, defined by transmission via efficient and selectively accessed paths [21]. In the broader context of network science, diffusion models are commonly used to describe spreading or cascading phenomena, such as the spread of news on social media platforms or epidemiological contagion in human contact networks [22– In contrast, routing protocols have found utility in systems such as postal, air-traffic and internet-infrastructure networks, where communication takes place via unique paths with a focus on efficiency and fidelity of information transfer [25– In neuroscience, these two conceptualizations of network communication strike opposing trade-offs in how they balance competing biological demands [14, 28]. Routing is thought to promote fast, faithful and frugal signalling by reducing the number of synapses traversed between communicating neural elements [13]. However, the identification of efficient routes is often predicated on global knowledge of network topology, a requirement unlikely to be met in decentralised biological systems [29]. Diffusion processes, conversely, do not require global information and could therefore be feasibility implemented in nervous systems, but are generally regarded as lossy and inefficient due to elevated numbers of signal retransmissions [21].

To date, most network modelling studies have considered that every pair of elements in the connectome communicates according to the same policy, e.g., via a unique model of routing or diffusion. This assumption presupposes a uniform mode of signalling across the brain and deems communication between all elements equally important. Connectomes, however, are heterogeneous networks, comprising elements with varied topological, spatial and neurobiological makeups [30–33]—factors that could engender different patterns of communication across the brain. For example, in the *Caenorhabditis elegans* connectome, different models may be suited to describe polysynaptic communication between distant sensory and motor neurons, as opposed to local interactions within a spatially compact and functionally homogeneous ganglion.

Here, we explore how connectome architecture may foster propensities towards diffusion or routing depending on the topological embedding of communicating node pairs. We focus on the well-documented modular organisation of connectomes [34–36], i.e., the tendency for neural elements to form assortative communities of dense internal connectivity and sparse external connectivity. This modular structure—observed across diverse connectivity modalities, spatiotemporal scales and species [37– 42]—is thought to provide a structural basis for functionally specialised systems. In the context of network communication, a module acts as a topological basin that concentrates or “traps” signal diffusion within its tight-knit connectivity, imposing structural barriers for signals to flow to other modules [43–46]. This observation tempers the notion that diffusion is universally inefficient, suggesting that connectome architecture may naturally steer passive spreading to achieve efficient communication between elements of the same module. Meanwhile, communication between elements belonging to different modules would benefit from a specialised routing policy capable of integrating information across long topological and spatial distances [47–50]. As such, it has been proposed that connectome communication may be preferentially modelled by diffusion within modules and routing between modules [5] (Fig 1A).

**FIG. 1.**
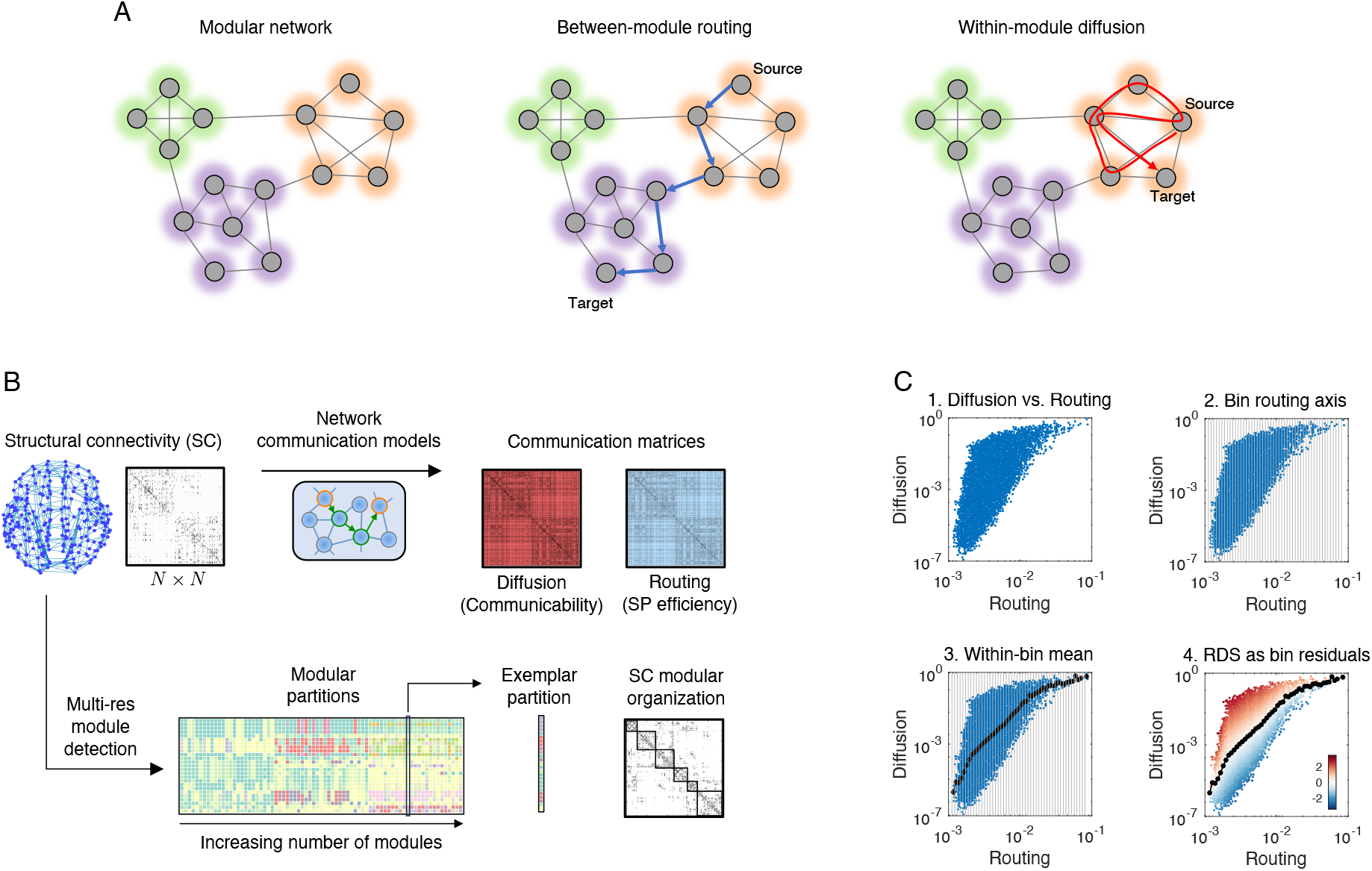
Methodology overview. **(A)** Left: Schematic of a modular network with 3 modules, shown in green, purple and orange. Each module is characterised by the presence of many internal connections and few external connections. Centre: Between-module communication may benefit from a selective path (blue) to efficiently access the rare connections bridging across modules. Right: Within-module communication may take advantage of dense internal connectivity to unfold via passive spreading (red), without the need for an efficient routing policy. **(B)** Top: Network communication models were used to estimate the structural capacity for signalling between every pair of nodes in a connectome. This yielded two communication matrices quantifying the capacity for diffusion (communicability) and routing (shortest path efficiency). Bottom: a routine based on the Louvain algorithm for modularity maximisation was used to compute the multi-resolution modular partitions of a connectome. **(C)** The relative diffusion score (RDS) was computed as the residual of the piecewise relationship between the diffusion and routing communication matrices. In brief, the routing axis of the routing-*vs*-diffusion scatter plot (1) was divided into equally spaced bins (2). Each bin was characterised by their mean and standard deviation capacity for diffusion (3). A node pair’s RDS was computed as their diffusion value minus the bin mean, divided by the bin standard deviation (4).

The aim of this paper is to investigate the hypothesis that connectome architecture favours within-module diffusion and between-module routing. To do this, we considered the connectomes of 6 organisms, chosen to cover a wide range of connectivity mapping techniques and spatial resolutions—from the complete nanoscale nervous systems of *C. elegans* to the inter-areal human connectome. We defined the relative diffusion score (RDS), a new measure to quantify the proportional capacity for neural communication via diffusion *vs* routing, and examine its interplay with connectome modular architecture. We posit that investigating whether connectome topology engenders heterogenous signalling regimes will advance our ability to faithfully model communication in complex brain networks.

## RESULTS

We analysed 6 previously published connectomes describing the structural connectivity (SC) of *C. elegans* [51], larval *Drosophila melanogaster* [19], adult *Drosophila melanogaster* [52], mouse [53], macaque [54] and human [55] brains (see *Methods*). For each connectome, we used network communication models to compute matrices quantifying the structural capacity for signalling between node pairs [56] (Fig 1B, top). Routing and diffusion were operationalized, respectively, via the *shortest path efficiency* and *communicability* measures. Shortest path efficiency quantifies communication via the least costly (most efficient) path between nodes, where cost refers to the number of connections, or the total connection length, of a path [57]. Routing via shortest paths is the most popular model of connectome communication [12, 58–60] and is at the core of many graph measures commonly used in network neuroscience (e.g., betweenness centrality, global efficiency, and the small-worldness index) [61]. Communicability conceptualises signalling as a broadcasting process that unfolds simultaneously along all walks in the network [62, 63]. In this model, communication capacity is not determined exclusively by the length of the shortest path, but also by the presence of multiple routes through which signals can travel between nodes. Communicability is one of the most widely used diffusion models in network neuroscience and the main alternative to shortest path-based measures [13], finding utility in a wide range of theoretical [64– experimental [17, 18] and clinical [67–69] research.

Brain network modules are not confined to a single organisational scale, instead displaying a multi-level and often hierarchical architecture [33, 34]. To account for this, the modular structure of each connectome was computed using a multi-resolution approach [70, 71] based on the Louvain algorithm for modularity maximisation [72, 73] (Fig 1B, bottom; *Methods*). For each connectome, this resulted in a rich ensemble of partitions ranging from coarse to fine decompositions. Throughout our analyses, we first focus on a partition with intermediary resolution obtained from this ensemble. We subsequently examine the entire set of partitions to characterise network communication as a function of modular organisational scales.

We developed the relative diffusion score (RDS) to quantify the proportional propensity for communication via routing or diffusion (Fig 1C; *Methods*). The RDS was inspired by conceptually similar benchmarking methodologies [74–76] and can be visualised in four steps. First, we plot the shortest path efficiency *vs* the communicability of each pair of nodes. Second, we divide the routing axis into equally spaced bins (vertical grey lines), such that node pairs within the same bin have approximately the same shortest path efficiency. Third, we compute the mean communicability within each bin (black dots). For each node pair, this value can be interpreted as their expected diffusion capacity given their routing capacity (i.e., the length of their topological shortest path). Lastly, the RDS is defined as the standardised residual with respect to the mean communicability within each bin. Node pairs with positive RDS (shown in red) show greater communicability than expected based on their shortest path efficiency, i.e. a propensity for diffusion over routing. Conversely, node pairs with negative RDS (shown in blue) show a propensity for routing over diffusion. Because RDS is defined as standardised residuals, values above 2 (below ™2) indicate significant (*α*=5%) propensities for diffusion (routing).

### Connectome architecture favours within-module diffusion and between-module routing

Figure 2 shows the SC, RDS and modular architecture of the 6 organisms considered in our paper. Connectomes varied in their size and connection density. The RDS is visualised in two formats: a routing-*vs*-diffusion scatter plot and a node-by-node matrix. SC and RDS matrices are ordered according to the modular partition of each connectome, such that node pairs clustered in the same module are positioned inside the main-diagonal blocks.

**FIG. 2.**
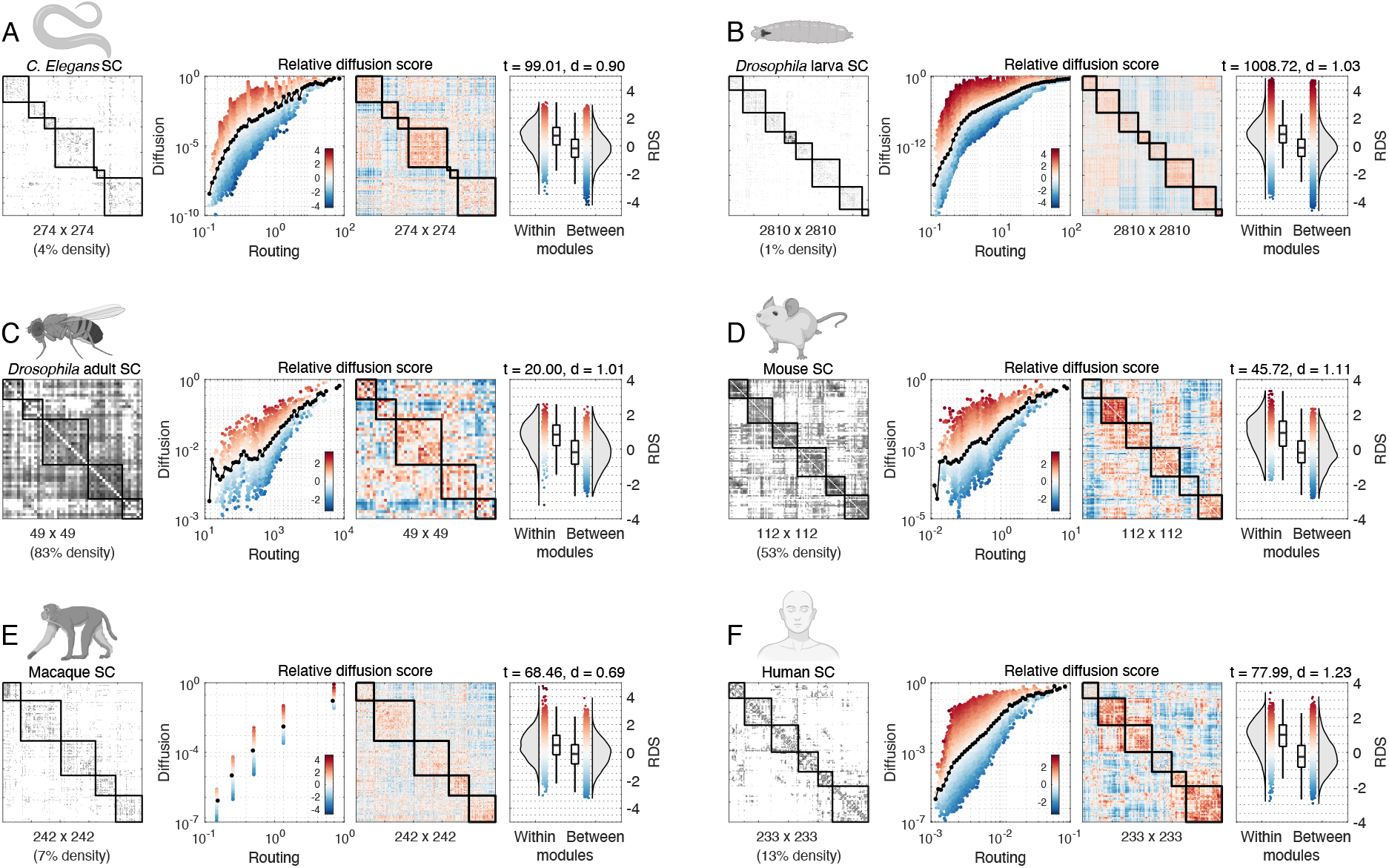
Interplay between structural connectivity, modular structure and RDS. From left to right, each panel shows the structural connectivity matrix, diffusion *vs* routing scatter plot, RDS matrix, and within- and between-module RDS distributions. Entries in the SC matrix are coloured according to connection weight. SC and RDS matrices are ordered such that nodes in the same module are positioned within the main-diagonal blocks. For each organism panel, RDS values are consistently coloured from red-to-blue across the scatter plot, matrix and boxplot visualisations. Boxplots report the *t*-statistic and Cohen’s *d* of a two-sample *t*-test comparing the means of within- and between-module RDS distributions.

Focusing on the routing-*vs*-diffusion scatter plots, a first observation is that shortest path efficiency and communicability are positively correlated for all organisms. This indicates that connectomes are wired such that, on average, node pairs with a high (low) capacity for routing also show a high (low) capacity for diffusion, despite the conceptual opposition between these models of communication [21]. Importantly, however, we did observe deviations from this mean trend, with certain pairs of nodes showing higher (RDS(*i, j*) *>* 0) or lower (RDS(*i, j*) *<* 0) capacity for diffusion than expected based on their capacity for routing.

We tested for differences in within- and between-module RDS distributions, and found that RDS is significantly larger within modules (two-sample *t*-tests, *p <* 10^*™*20^ for all connectomes). Although *t*-statistics varied as a function of network size (connectomes with more nodes yielded larger *t*-statistics due to more degrees of freedom), the Cohen’s *d* effect size was comparable across organisms (average *d*=0.99, ranging from 0.69 for the macaque to 1.23 for the human connectomes). Therefore, despite marked differences in connectome size, density and spatial scale, we observed a strong and stable effect of modular structure on RDS, pointing towards a propensity for diffusion within modules and routing between modules.

Building on these initial findings, we tested if RDS alone could classify whether nodes belong to the same or different modules. Casting the analysis in terms of a classification problem moves beyond simply detecting statistical differences and towards predictions about connectome architecture. To do this, we first selected node pairs with |RDS(*i, j*)| *>* 2, i.e., those with statistically significant deviations towards routing or diffusion at *α*=5%. Node pairs with RDS(*i, j*) *>* 2 (diffusion-prone) and RDS(*i, j*) *< ™*2 (routing-prone) were classified, respectively, as within- and between-module pairs. Figure 3 shows that this simple classification scheme was able to retrieve modular co-assignments with high accuracy (average accuracy of 83.5%, ranging from 76% for the human and 93% for the mouse connectomes). Routing-prone node pairs very often belonged to different modules (in 97.2% of cases on average; RDS *<* ™2 columns in the confusion matrices), while diffusion-prone pairs, although predominantly positioned within the same module (70.3% on average; RDS *>* 2 columns), were relatively harder to classify. Conversely, almost all within-module node pairs with significant RDS were prone to diffusion (96.0% on average; *Within* rows), while a relatively lower fraction of between-module pairs were routing-prone (75.6% on average; *Between* rows).

**FIG. 3.**
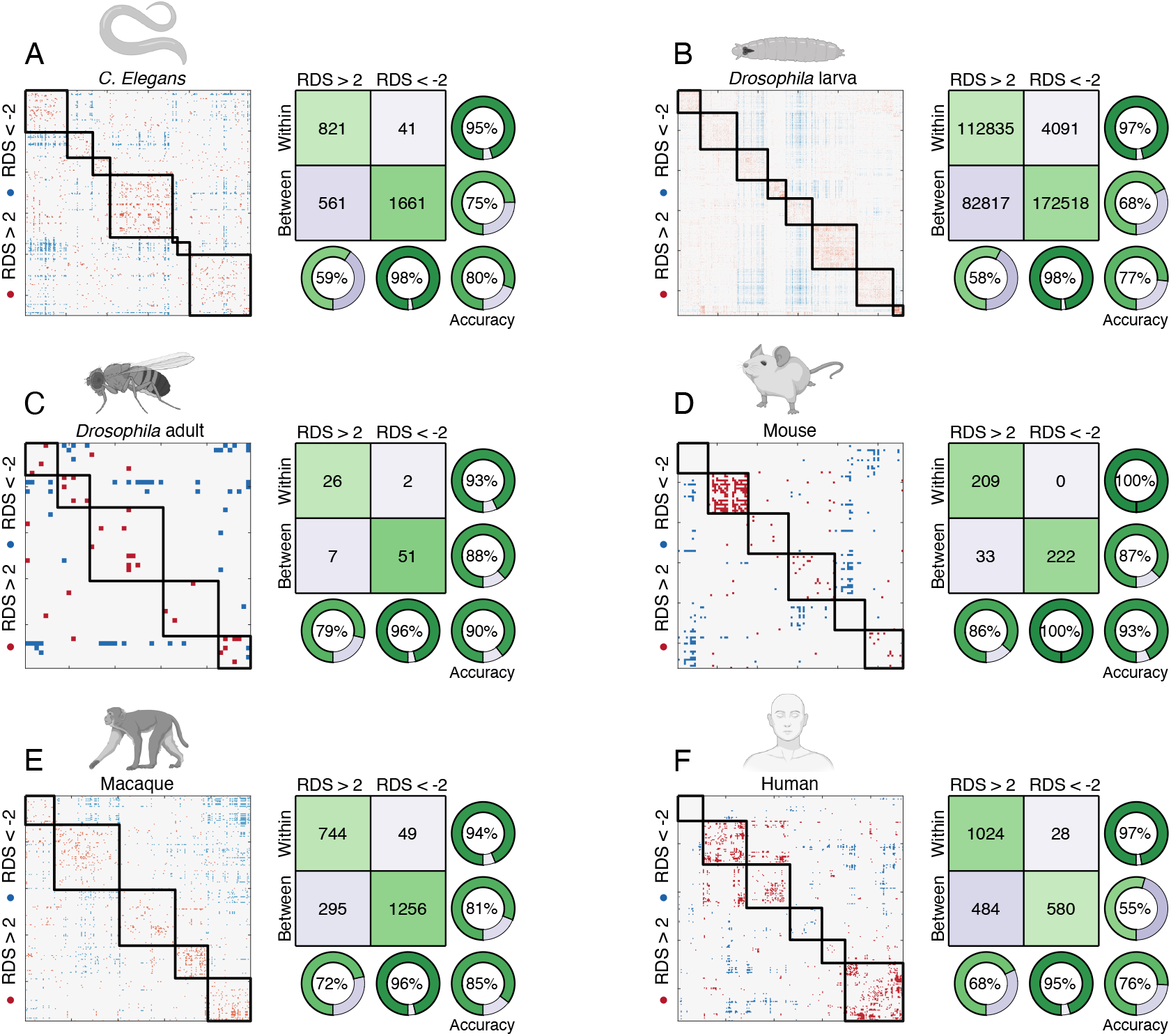
Prediction of modular co-assignment using RDS. RDS matrices were thresholded to show only significant propensities towards diffusion (RDS *>* 2, red entries) or routing (RDS *<* − 2, blue entries). Node pairs with red and blue entries were classified as within- and between-modules, respectively. Confusion matrices visualise the performance of this classification scheme, with the actual modular co-assignment shown in the rows (labels Within and Between) and the RDS prediction shown in the columns (labels RDS *>* 2 and RDS *<* −2). Pie charts show the proportion of correctly classified node pairs along different dimensions of the confusion matrix and the overall classification accuracy.

Taken together, this first set of results indicates a structured interplay between the RDS and the modular architecture of connectomes. Specifically, RDS was (i) significantly larger within modules than between modules and (ii) predictive of whether two nodes are assigned to the same or different modules. The direction of these associations—which was consistent across drastically different species and spatial scales—was such that intra- and inter-module communication showed a propensity towards diffusion and routing, respectively. This finding supports the hypothesis that brain network communication may be organised to support within-module diffusion and between-module routing.

### Topological and geometric influences on the interplay between routing, diffusion and modular architecture

What factors account for the observed relationship between network communication and modular architecture? In this section, we investigated a number of topological and geometric connectome properties as putative determinants for the propensity towards diffusion or routing. As a starting point for these analyses, we draw the reader’s attention back to Figure Fig 2. While we have seen that, on average, RDS is greater within modules, examination of the RDS matrices reveals that this tendency is not homogeneous across modules. For example, this is clear in the mouse connectome (Fig 2D), where the second block along the main diagonal shows visibly larger RDS compared to other modules. What accounts for this heterogeneity?

To examine this question, in Figure 4, we considered the average RDS of each modular block, i.e., within each module and between every pair of modules. This condenses the node-by-node RDS matrix into a module-by-module block representation. Next, we computed the connection density (number of connections divided by number of node pairs) and the spatial compactness (average Euclidean Distance between nodes) of every modular block. Examining the associations between these measures and average RDS, we found that denser and more spatially compact blocks showed a stronger propensity towards diffusion, while sparser and more spatially dispersed blocks were more prone to routing—an observation that was once again consistent in all connectomes.

**FIG. 4.**
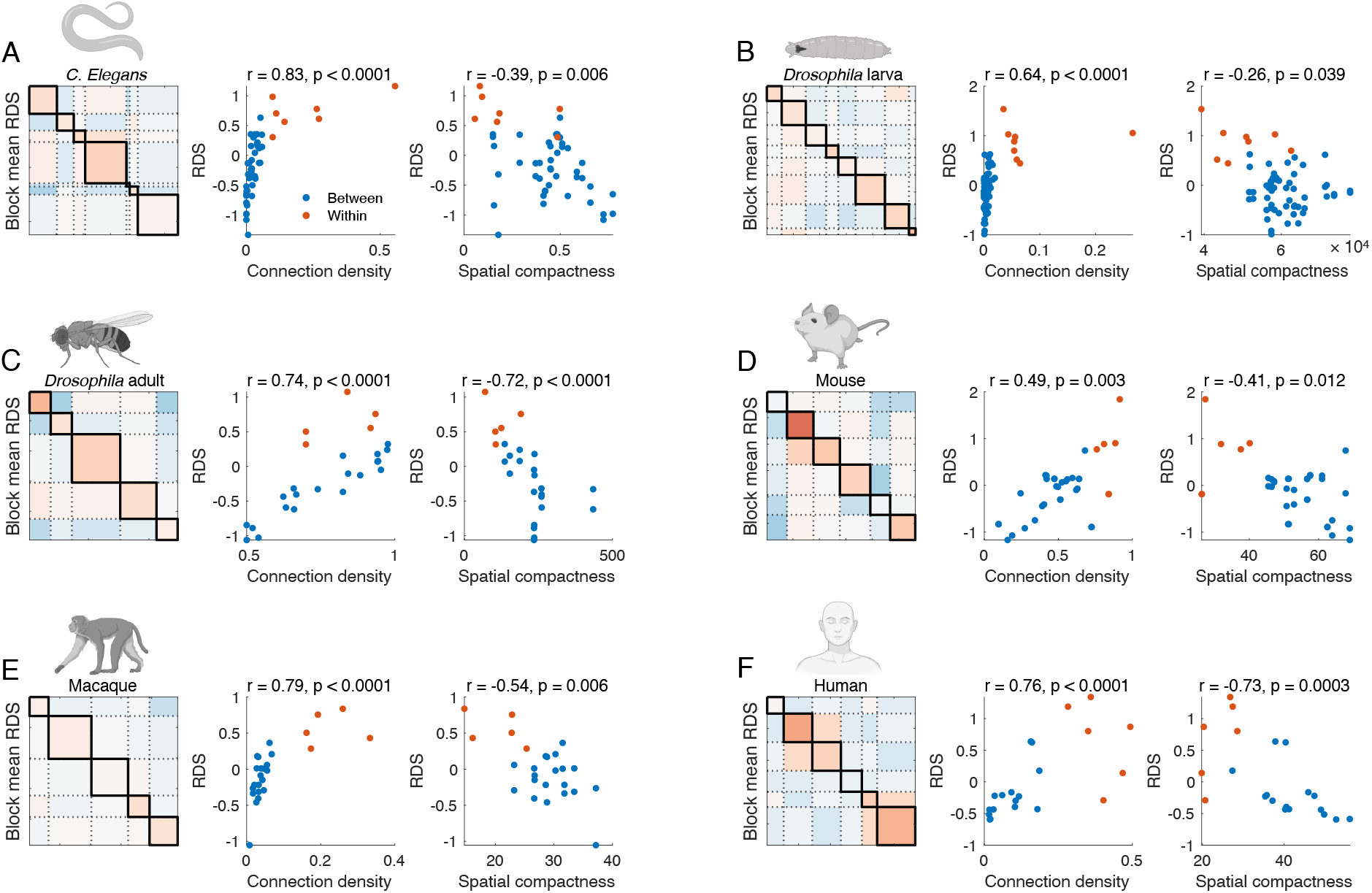
Relationship between RDS, connection density and spatial compactness in modular blocks. A modular block can refer to nodes within the same module (main-diagonal blocks, red dots) or between two different modules (off-diagonal blocks, blue dots). The panel of each organism shows their RDS matrix downsampled to the level of modular blocks, such that each block is characterised by the mean RDS of its node pairs. Scatter plots show the association between block RDS and (left) block connection density and (right) block spatial compactness, as measured by the Spearman rank correlation.

We note that, by design, nodes in the same module tend to be more densely connected to each other than to the rest of the network [77]. In addition, given the well-documented relationship between neural connectivity and spatial distance [78, 79], we also expect nodes within the same module to be in closer proximity. Both of these expectations were met in our analyses. What is interesting here is that the natural heterogeneity in these expectations (i.e., some modular blocks will happen to be denser or more compact than others) was correlated to the heterogeneity in the block RDS. This result echoes the intuition behind our paper’s central hypothesis: diffusion, predicated on passive spreading dynamics, may be more suited to describe localised communication between tight-knit clusters of neural elements, while routing, facilitated by selective and efficient paths, may be a better model of long-range signalling across segregated brain systems.

We sought to further untangle the relationship between modular architecture, spatial proximity, and network communication by means of null network models [80]. Ensembles of null networks were constructed by randomly rewiring connectomes while preserving topological and geometric properties of interest. We considered three null network models: (i) *Rewired*, where connection were randomly rewired while preserving the degree sequence of original connectomes [81]; (ii) *Cost-preserving rewired*, where the rewiring additionally preserved the distribution of connection lengths and the relationship between connection weights and lengths [82]; and (iii) *Block rewired*, where the rewiring again preserved degree sequence, but was additionally restricted to take place within modular blocks, thus conversing the modular architecture of original connectomes. Repeating our analyses in these null networks allowed us to probe which connectome features contribute to the observed dynamics between diffusion and routing. For each null network in an ensemble, we tested for statistical differences between within- and between-module RDS distributions. Each ensemble comprised 1,000 networks, yielding a null distribution of *t*-statistics that we compared to the results obtained in the original connectomes.

As shown in Figure 5, complete rewiring of network topology eliminated the interplay between modular architecture and RDS (*Rewired* null model, purple histograms). This is unsurprising, as indiscriminate rewiring dismantles the modular structure of original connectomes. In contrast, rewiring while preserving connection lengths yielded null networks that recapitulated the original concentration of high RDS within modules (*Cost-preserving rewired* null model, green histograms). While the strength of this association did not match that of empirical connectomes (dashed vertical lines), it nevertheless underscores the well-established relationship between modules and distance, indicating that the spatial embedding of brain networks partially explains the interplay between diffusion, routing and modules (see also Fig S1). Interestingly, preserving the original modular architecture of connectomes while otherwise randomising their topology led to an increase in the propensity for within-module diffusion and between-module routing (*Block rewired* null model, yellow histograms). This indicates that connectome topology does not maximise this propensity, and instead may have other features that contribute to the relative capacity for diffusion and routing. For example, block-rewired networks are expected to have longer shortest-path lengths and smaller clustering coefficient within each modular block compared to empirical connectomes, properties that have been reported to influence network communication [27].

**FIG. 5.**
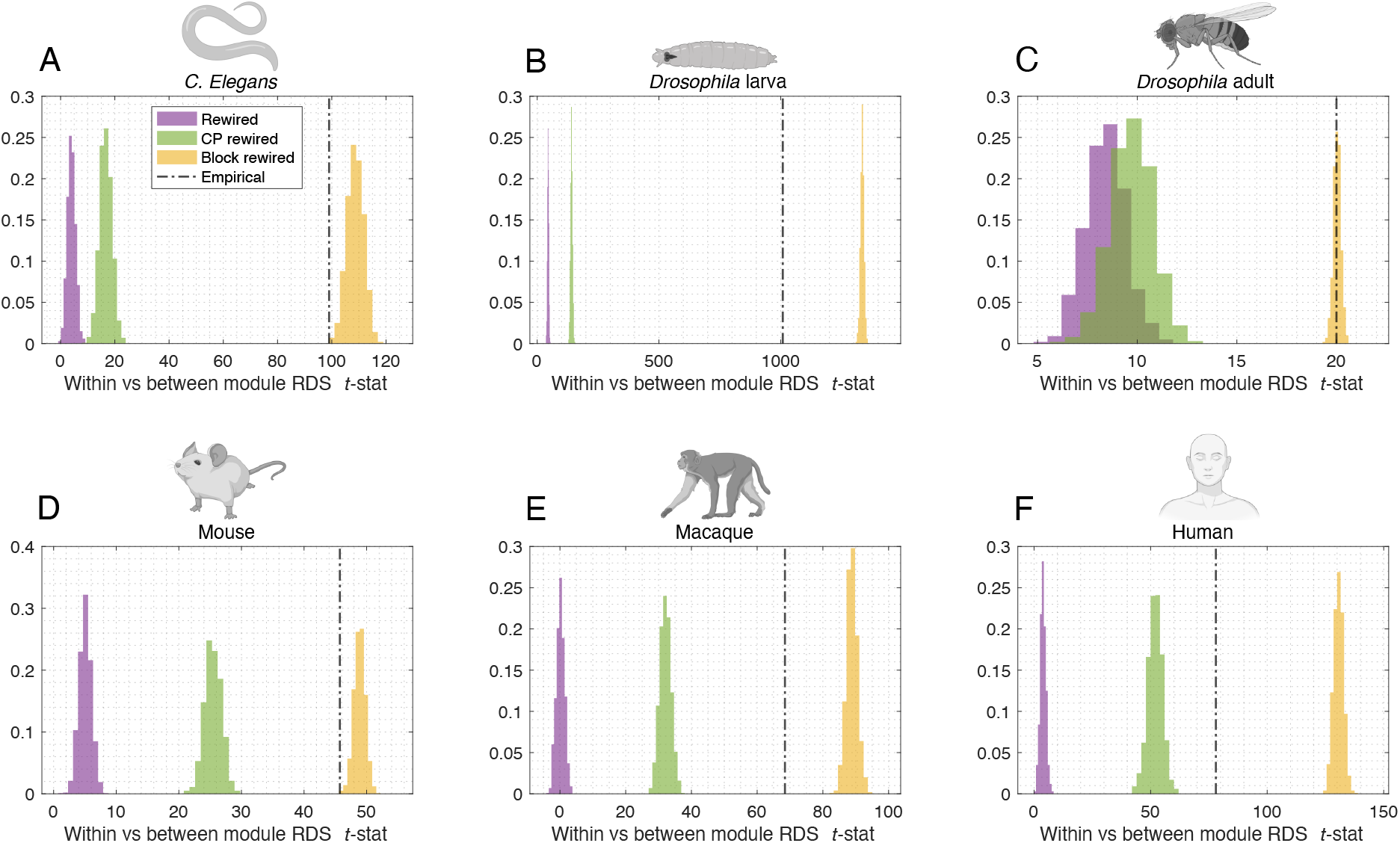
RDS in null network models. Histograms show the distributions of *t*-statistics from comparing within-*vs* between-module RDS distributions in ensembles of 1,000 *Rewired* (purple), *Cost-preserving rewired* (green), and *Block rewired* (yellow) null networks. Dashed vertical lines show the *t*-statistics from original connectomes.

Collectively, the results of this section shed light onto the observed propensity for within-module diffusion and between-module routing in connectomes. We found that heterogeneity in this mean trend is explained by inter- and intra-module connection density and spatial proximity. Accordingly, our rewiring analysis suggests that the interplay between diffusion, routing and modular structure is contingent on the fine-tuned topological and geometric organisation of connectomes.

### Interplay between routing, diffusion and multi-resolution modular architecture

So far, our analyses have been based on a single modular partition for each connectome. While this approach is useful due to its tractability, it overlooks two important facets of brain network organisation. First, it presupposes a fixed number of modules for each network, thus focusing on a potentially arbitrary modular resolution that may not reflect the multi-level organisation of neural connectivity. Second, it disregards the degeneracy of modular structure. This means that, for a fixed number of modules, different partitions can decompose a network into modules with the same or very similar aptitude—a property that is reflected in the stochasticity inherent to many algorithms for community detection [83].

In this section, we investigate the impact of modular resolution and degeneracy in the interplay between diffusion and routing. To do this, we examined increasingly fine partitions, ranging from 2 to up to 100 modules, depending on the organism. In brief, multi-resolution partition sets were computed by systematically varying the resolution parameter *γ*, which governs the number of modules identified by the Louvain algorithm (Fig 1B, bottom; see *Methods*). For each value of *γ*, we computed 100 independent partitions.

Figure 6 shows the *t*-statistics comparing within-*vs* between-module RDS, as a function of modular resolution. Each grey dot marks the *t*-statistic and the number of modules of a single partition, while the black line tracks the average statistic. First, we note that this average is always positive, meaning that the propensity for within-module diffusion and between-module routing is evident for partitions of all granularities. Interestingly, the *t*-statistics peaked at a particular resolution, which was similar across organisms (6 modules on average, ranging from 3 in the *C. elegans* to 10 in the mouse connectomes; dashed green lines). In most cases, these peaks were not aligned with the partition considered in previous analyses (dashed purple lines), demonstrating that our previous results did not focus on the partitions that maximised the support for our hypothesis. In addition, we found that different partitions with the same number of modules led to consistent *t*-statistics, indicating that our results are robust to the stochasticity of the Louvain algorithm.

**FIG. 6.**
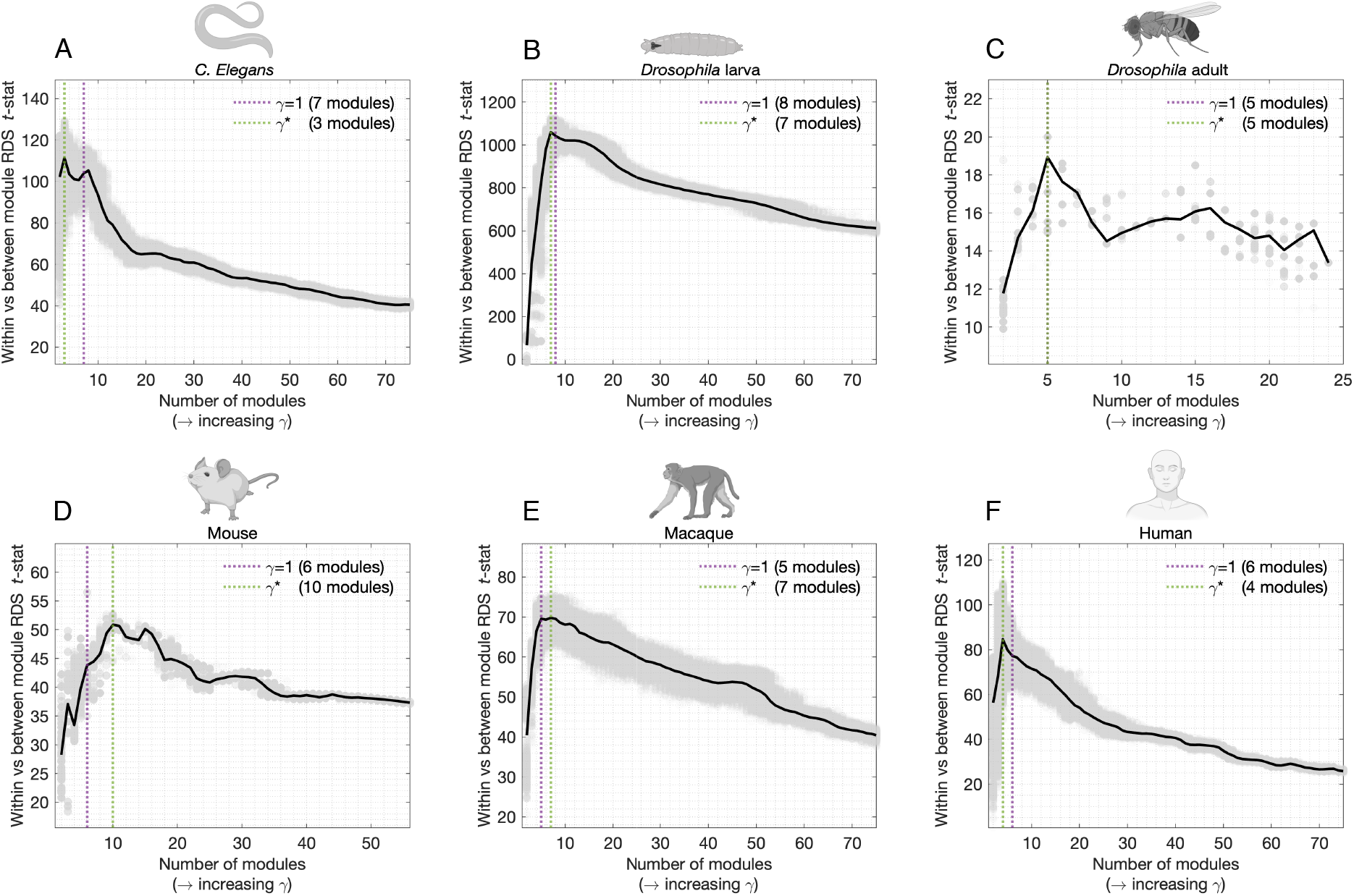
Interplay RDS and multi-resolution modular structure. We computed partitions with increasing number of modules by varying the resolution parameter *γ*. A set of 100 partitions were computed for each value of *γ* to account for the stochastic behaviour of the Louvain algorithm. In each organism panel, a grey dot denotes a single partition, characterised by its number of modules and the *t*-statistic from the within-*vs* between-module RDS comparison. The black curve tracks the average *t*-statistic across partition resolutions. For ease of visualisation, we consider partitions ranging from 2 to 75 modules (with the expectation of the adult fly, for which the connectome comprises only 49 nodes and the *x*-axis was capped at 25 modules). Dashed vertical lines mark the partition from previous analyses (purple) and the partitions with the largest mean *t*-statistic (green).

The peaks in the *t*-statistic curves can be understood as follows. If a partition is too coarse, modules will be large enough to encompass node pairs separated by long topological distances (e.g., many hops along the shortest path). Such node pairs will tend to have low diffusion capacity and therefore their assignment to the same module will hamper within-module diffusion. Conversely, if a partition is too fine, node pairs that are topologically close will be split into different modules, therefore increasing the capacity for between-module diffusion. Both of these scenarios will contribute to decreasing the *t*-statistic curves in Figure 6, thus resulting in narrow peaks at intermediate partition resolutions. This finding suggests the existence of a connectome organisational scale—characterised by approximately 5 to 7 modules— that amplifies the contrast between inter-module diffusion and intra-module routing. We conjecture that this architectural scale may offer a sweet spot to study the communication dynamics of brain networks of different species and spatial dimensions.

#### Sensitivity analyses

Lastly, we close our analyses by testing the sensitivity of our results to a range of methodological choices and parameters. In Figure S2, we revisited the classification problem of Figure 3 to investigate the prediction accuracy of modular co-assignment as a function of partition resolution. In close agreement with the results from the previous section, we found that prediction accuracies peaked around partitions with 5 to 7 modules and were consistent across independent runs of the modular decomposition algorithm. In Figure S3, we investigated how the effect size of within-*vs* between-module RDS was affected by (i) the number of *x*-axis bins in RDS computation, (ii) the use non-parametric statistical tests, (iii) rank-transformations of network communication measures prior to RDS computation, and (iv) connection directionality. Our results remained consistent across all explored scenarios.

## DISCUSSION

In this study, we investigated the hypothesis that connectome communication is preferentially modelled by diffusion within modules and routing between modules. To test this, we introduced the relative diffusion score (RDS), a novel measure that benchmarks the structural capacity of node pairs to communicate via diffusion *versus* routing. By examining the relationship between RDS and the modular architecture of connectomes across six organisms, we found multiple lines of evidence supporting the central hypothesis. These findings, consistent across species with diverse connectome mapping techniques, spatial resolutions, and connectivity densities, suggest that communication in brain networks may favour a hybrid signalling strategy—passive diffusion within modules and selective routing between modules.

### Brain network communication and modular structure

Modularity is one of the most fundamental features of connectomes. In humans, structural modules are the anatomical basis for functionally specialised systems [36, 71] conjectured as the building blocks of higher-order cognition [84, 85]. Indeed, the interplay between intramodule segregation and inter-module integration is at the core of several theories of neurodevelopment [86, 87] and cognition [88] as network-based processes. Relatedly, recent works at the intersection of neuroscience and machine learning indicate that modularity is critical for dynamic computation [89–91] and a key emergent property of biologically constrained artificial neural networks [66, 92]. Our results add to the growing literature on the importance of modularity in nervous systems, and posit a new role for this architecture as a mediator of hybrid modes of connectome communication. The finding that within-module diffusion is more efficient than expected suggests that passive spreading dynamics could support spatially localised and functionally segregated information processing, without the need for global knowledge of the network to guide signalling. Routing via selective and efficient polysynaptic paths might be then reserved to bridge across neural elements embedded in different functional systems or separated by long topological distances, for which diffusion would entail elevated delays and numbers of signal retransmission.

The conceptualisation of neural communication as a hybrid diffusion-routing process provides new perspectives on the study of brain network economy, i.e., the trade-off between wiring cost and adaptive functionality of connectome topology [48]. Hubs, rich clubs, and longrange connections are expensive features to maintain and develop [58]. The architectural investment in these properties is thought justified by their role in a global backbone of shortest paths, which enables polysynaptic communication between any two parts of the brain in a small number of steps [47, 49]. Concurrently, connectomes are embedded in spatial, hyperbolic and functional metric spaces that facilitate the use of hubs and rich clubs for the decentralised identification of efficient paths [50, 93–95]. Based on these points, it would seem that connectome architecture is organised to promote routing, and that diffusion would not take advantage of the investment in costly topological features. However, the assumption of routing as the sole communication policy also has important limitations. For example, connection weights follow a heterogeneous distribution, featuring many weak connections and rare strong ones [13]. Empirical analyses of the human connectome show that shortest paths only use a small subset of high-weight connections, with the majority of white matter tracts in the brain not involved in any communication via shortest paths [50, 96]. At the same time, hubs and strong connections are overburdened, forming traffic bottlenecks that could contribute to signalling delays and losses in fidelity [97–99]. Considering a composite of diffusion and routing mediated by modular boundaries conciliates the pros and cons of both strategies. Diffusion percolates via low-weight connections within modules, which contribute to functional compartmentalisation by concentrating information flow to localised circuits of dense connectivity. Meanwhile, dedicated routing processes utilise the connectome’s expensive backbone to promote efficient, long-range and cross-system functional integration.

An open challenge in the study of modularity is that complex networks are seldom characterised by a single organisational scale. For instance, in the case of the human brain, the foremost modular decomposition of cortical regions into functional systems is a multi-level partition into 7 and 17 modules [85]. In most cases, there is little consensus as to what level of modular granularity should be prioritised [83], especially when dealing with connec-tomes from different species and spatial scales. While recent methods now allow for the tractable characterisation of the hierarchical structure of a network [70, 100], there remains the question of which modular resolution should be prioritised when studying brain dynamics. By examining the interplay between diffusion and routing across modular granularities, our findings suggest an intermediate scale—around 5 to 7 modules—as a putative universal resolution to balance segregation and integration in connectomes.

### Future directions

Several technical aspects of our work could be further investigated in future research. First, we operationalise routing and diffusion using shortest path efficiency and communicability, but other network communication measures such as navigation [50], mean first passage time [21], or search information [29] could be explored. Second, the piecewise linear method used to define RDS could be replaced by continuous methods based on spline regression or exponential fits, which would depend on more statistical parameters but bypass the need for dividing data points into bins. Third, methods such as stochastic block models provide an alternative to community detection via modularity maximisation, which could be used to explore network communication in non-assortativity modular organisations [100–102]. Finally, instead of modules computed from structural connectomes, it would be interesting to consider the interplay between RDS and functional modules, such as human canonical resting-state networks [85] or communities of neurons associated with circumscribed functional roles in *C. elegans* or *Drosophila* [33].

More broadly, our focus on modularity is only one out of many possible approaches to model hybrid connectome communication. Other hallmark properties of brain networks, such as homophily, hubness or rich-club allegiance, could be used to adjudicate the selection of different signalling policies. Beyond network topology, the increasing availability of rich annotations on the neural elements comprising connectomes—from macroscale human brain regions to individual neurons in model organisms—potentiates the use of morphological [19], molecular [103], cytoarchitectonic [104], genetic [105], and dynamical [106] information to enrich network models. Early evidence suggests that inter-areal variation along some of these properties may reflect propensities towards different signalling strategies [94, 107–109]. In parallel, other reports have proposed that the explanatory power of different communication models may vary according to behavioural and cognitive contexts [56, 110], as well as to species and brain volume [37, 111]. Untangling these observations will require theoretical efforts to develop communication models that explicitly incorporate different types of annotations [30, 31]. Such annotated network measures would extend the simple approach used in this paper, where modules act as hard boundaries dictating the switch between routing and diffusion. For example, recent advances in the formalisation of biased random walks enable the flexible interpolation of composite communication models using an arbitrary nodal attribute [28, 112–114], providing a promising avenue of future exploration.

We envision that progress in the research directions above will approximate network communication models to mechanistic accounts of brain function. While successful in explaining a range of experimental observations— from electrical pulse propagation [17] to the functional effects of targeted lesioning [18]—the present class of models is not explicitly grounded in neuronal physiology. Model parametrization using relevant annotations stands to narrow this gap, for example by revealing biophysical or neurochemical cues that could support dynamic switching between modes of communication. Importantly, empirical validation will be critical to identify models and annotations faithful to biological neural signalling [14], e.g., by using maps of causal stimulus transmission [60, 115] or methods to infer communication from neural recordings [116, 117]. In turn, we posit that advances in our understanding of connectome communication will inform related computational domains predicated on simulating neural interactions, potentiating the design of dynamical and generative models of the brain [64, 66, 118–120].

## METHODS

### Connectivity data

A connectome comprising *n* nodes is represented by a connectivity matrix 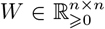. The entry *W*_*ij*_ denotes the strength of the structural connection from node *I* to node *j*, where *W*_*ij*_ = 0 indicates the absence of a connection. We considered 6 previously published and openly available connectomes from a range of organisms, as detailed below. Brain networks were weighted (except the macaque connectome), with weights reflecting the strength of physical links between neural elements, and directed (except the human connectome), with connections capturing the orientation of information flow. We note that modelling network communication requires connectomes to be strongly connected, i.e., each node must have at least one incoming and one outgoing connection. When this requirement was not met by the original connectomes, we considered the subset of nodes comprising the network’s strongly connected component.

#### Caenorhabditis elegans

Complete nanoscale connectome comprising electrical and chemical synapses between 302 neurons mapped using electron microscopy [2, 121, 122]. Connections are directed and weights denote the number of synaptic contacts between pre- and postsynaptic neurons. Following [43], we considered the connectivity between 279 neurons of the somatic nervous systems (20 pharyngeal and 3 somatic neurons that do not form connections to the rest of the network were excluded) with 3.9% connection density.

#### Larval Drosophila melanogaster

Complete nanoscale connectome comprising axon-axon, axon-dendrite, dendrite-axon, and dendrite-dendrite synaptic connectivity between 3,016 neurons, reconstructed using electron microscopy and machine learning [19]. Connections are directed and weights denote the number of synaptic contacts between pre- and postsynaptic neurons. We considered the original connectome’s strongly connected component formed by a subset of 2,810 neurons with 1.3% connection density.

#### Adult Drosophila melanogaster

Connectome describing neural projections between 49 macroscale regions (local processing units) of the adult fly brain [52]. Connections are directed and weights denote the strength of projections between regions, inferred by aggregating single-cell resolution images from the FlyCircuit database [123]. The original connectome from [52] is strongly connected and has 83% connection density.

#### Mouse

Connectome describing mesoscale connectivity between 112 cortical and subcortical regions of the mouse brain [53], mapped using anterograde tract-tracing data from The Allen Brain Institute [3]. Connections represent directed interregional axonal projections and their weights were determined as the proportion of tracer density found in target and injected regions. The original connectome from [53] is strongly connected and has 53% connection density.

#### Macaque

Macroscale connectome comprising 383 cortical and subcortical regions of the macaque brain [54]. Connectivity data was collated from 410 anatomical tracing studies compiled in the CoCo-Mac database [124]. Connections represent directed long-distance tracts and are binary, indicating either the presence or absence of at least one study in the database documenting a tract between two regions. We considered the original connectome’s strongly connected component formed by a subset of 242 regions with 7.0% connection density.

#### Human

Macroscale connectome describing connectivity between 219 cortical and 14 subcortical regions from [55, 125]. Connectomes were constructed for 70 healthy adult participants using diffusion weighted imaging and deterministic white matter tractography. Connections are undirected and their weights are a proxy of the white matter fibre density between regions [126]. Following [125], subjects’ connectivity matrices were combined into a group-consensus connectome, which mitigates idiosyncratic reconstruction inaccuracies while preserving key features of individual brain networks. The original connectome from [125] is strongly connected and has 12.6% connection density.

### Network communication models

#### Shortest path efficiency

The identification of shortest paths is a minimisation problem that requires the remapping of connection weights (a measure of affinity, strength, or ease of interaction between nodes) into connection lengths (difficulty, delay or cost of interaction). We applied the transformation *L* = 1*/W*, which performs a monotonic remapping of strong weights into short lengths. The shortest path length from node *i* to *j* is 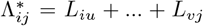, where {*u*, …, *v*} are the intermediate nodes along the shortest path. The shortest path efficiency was computed as 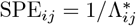[57].

#### Communicability

Communicability models neural signalling as a diffusive process that broadcasts information along all possible walks in the network. This notion is formalised as a weighted sum of the total number of walks between two nodes, where the weight of each walk is inversely proportional to their length. The computation of communicability is typically preceded by a normalisation of the entries of the connectivity matrix as 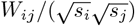, where *s*_*i*_ is the strength of node *i* [69]. This step aims to attenuate the influence of high strength nodes in the communication dynamics. Here, to account for the directed nature of the connectomes under examination, we generalised this normalisation as 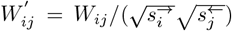, where right and left arrows indicate out- and in-strength, respectively. The communicability from node *i* to node *j* was then computed as 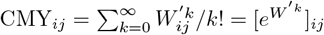 [62].

### Relative diffusion score

Prior to the computation of RDS, the shortest path efficiency and communicability matrices were logtransformed (base 10) to account for their heterogeneous nature, yielding approximately log-normal distributions of routing and diffusion capacity between nodes. As an alternative, we also considered a rank-transformation of communication matrices (see Fig S3). The range of shortest path efficiency values was divided into 50 equally sized bins (also see Fig S3 for alternative numbers of bins), such that each pair of nodes was assigned to a single bin *b*. Let *B*_*b*_ be the set of nodes in *b*. The RDS of node pair *ij* in bin *b* was defined as 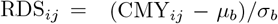, where 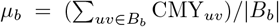 and 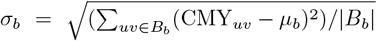 . In other words, the RDS is standardised residual with respect to the mean communicability within each shortest path efficiency bin. It quantifies deviations from a node pair’s expected capacity for diffusion given its capacity for routing.

We note that the choice of binning the routing axis (instead of the diffusion one) was motivated from the fact that the topological distance via the shortest path remains the mainstay measure of communication in brain networks [14]. As such, the RDS quantifies the capacity for diffusion relative to the well-characterised measure of shortest path efficiency. Alternatively, one could bin the diffusion axis to define the “relative routing score”, which would benchmark routing in relation to diffusion.

### Multi-resolution community detection

Structural connectivity modules were delineated using the Louvain algorithm for modularity maximisation [72, 77]. The number of modules identified by this method depends on the resolution parameter *γ*. For a given connectivity matrix *W* and *γ*, the Louvain algorithm seeks to maximise the modularity statistic *Q* = Σ_*ij*_ (*W*_*ij*_ ™ *γB*_*ij*_)*δ*_*ij*_, where *δ*_*ij*_ = 1 if *i* and *j* are assigned to the same module and *δ*_*ij*_ = 0 otherwise, and *B*_*ij*_ is the expected weight of the *ij* connection under a null model. Following previous work on community detection in structural connectomes [37, 39], we used the Girvan-Newman null model *B*_*ij*_ = *s*_*i*_*s*_*j*_*/*2*m* [127], where *s*_*i*_ is the strength of node *i* and *m* the number of connections in the network.

To account for the multi-scale organisation of brain networks, we used a multi-resolution routine based on a double sweep on the *γ* parameter space [70, 71]. In the first pass of the routine, we considered a sample of 1,000 logarithmically spaced *γ* values, ranging from 0.001 to 100. The first sweep was used to identify *γ*_*p*_ and *γ*_*q*_, parameters that led to, respectively, the first partition with at least 2 modules and the last partition with no more than min(*n/*2, 100) modules. The upper limit of 100 modules was set to ensure computational tractability when dealing with large connectivity matrices (e.g., *n* = 2810 for the larval fly). In the second pass of the routine, we sampled a new set of 1,000 logarithmically spaced parameters within the narrower range from *γ*_*p*_ to *γ*_*q*_. This ensures the construction of a densely sampled partition ensemble confined within biologically realistic module resolutions. Finally, to account for the stochasticity of the Louvain algorithm, we repeat the routine above 100 times, resulting, for which connectome, in a 100 *×* 1,000 multi-resolution partition ensemble. Figs 2 to 5 analyse an intermediate-resolution partition from this ensemble, defined as the one with median *Q* across the 100 repetitions for the default parameter *γ* = 1. Fig 6 analyses the entire ensemble of partitions.

## Data and code availability

Connectomes are publicly available through the references of section *Connectivity data*. Network communication models were computed using the Brain Connectivity Toolbox [61]. The routine for multi-resolution community detection was implemented by adapting code made available in [70]. Rewiring null models were implemented using code from the Brain Connectivity Toolbox [61] and [82].

---

**FIG. S1.**
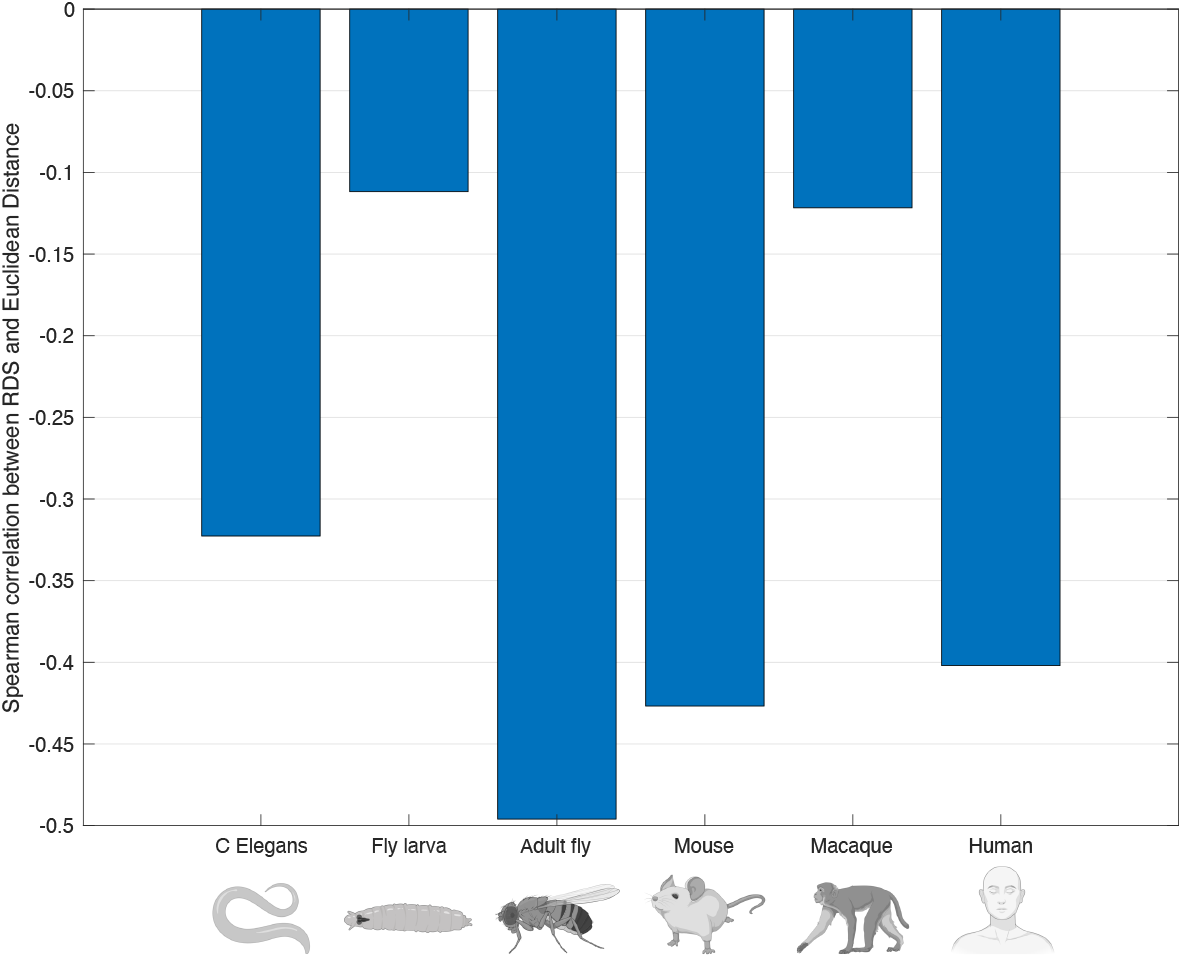
Spearman rank correlation between inter-element Euclidean distance and RDS.

**FIG. S2.**
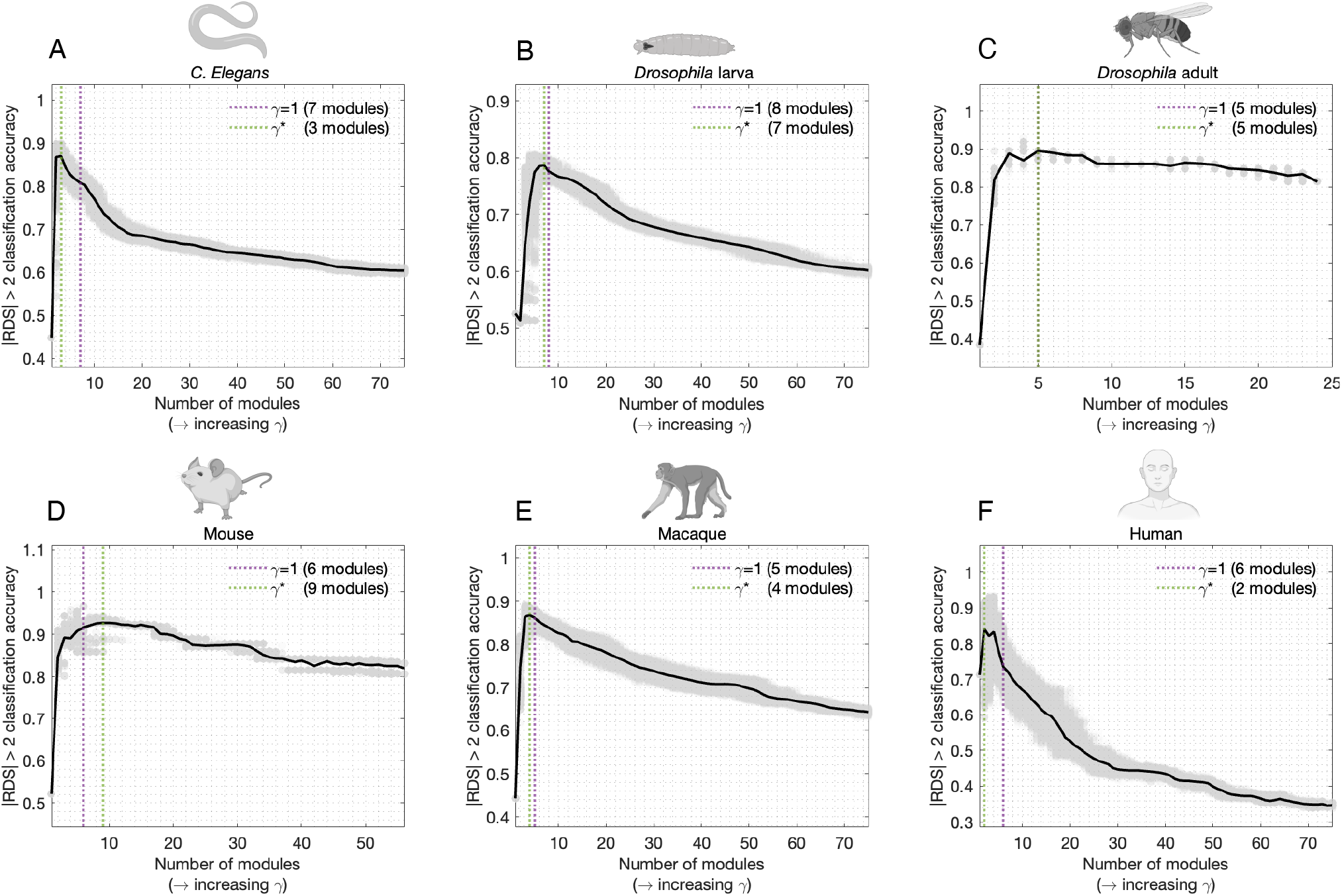
Prediction of community co-assignment using RDS across modular resolutions. We computed partitions with increasing number of modules by varying the resolution parameter *γ*. A set of 100 partitions were computed for each value of *γ* to account for the stochastic behaviour of the Louvain algorithm. Node pairs with RDS *>* 2 and RDS *< −*2 entries were classified as within- and between-modules, respectively. In each organism panel, a grey dot denotes a single partition, characterised by its number of modules and classification accuracy. The black curve tracks the average accuracy across partition resolutions. For ease of visualisation, we consider partitions ranging from 2 to 75 modules (with the expectation of the adult fly, for which the connectome comprises only 49 nodes and the *x*-axis was capped at 25 modules). Dashed vertical lines mark the partition from previous analyses (purple) and the partitions with the greatest mean accuracy (green).

**FIG. S3.**
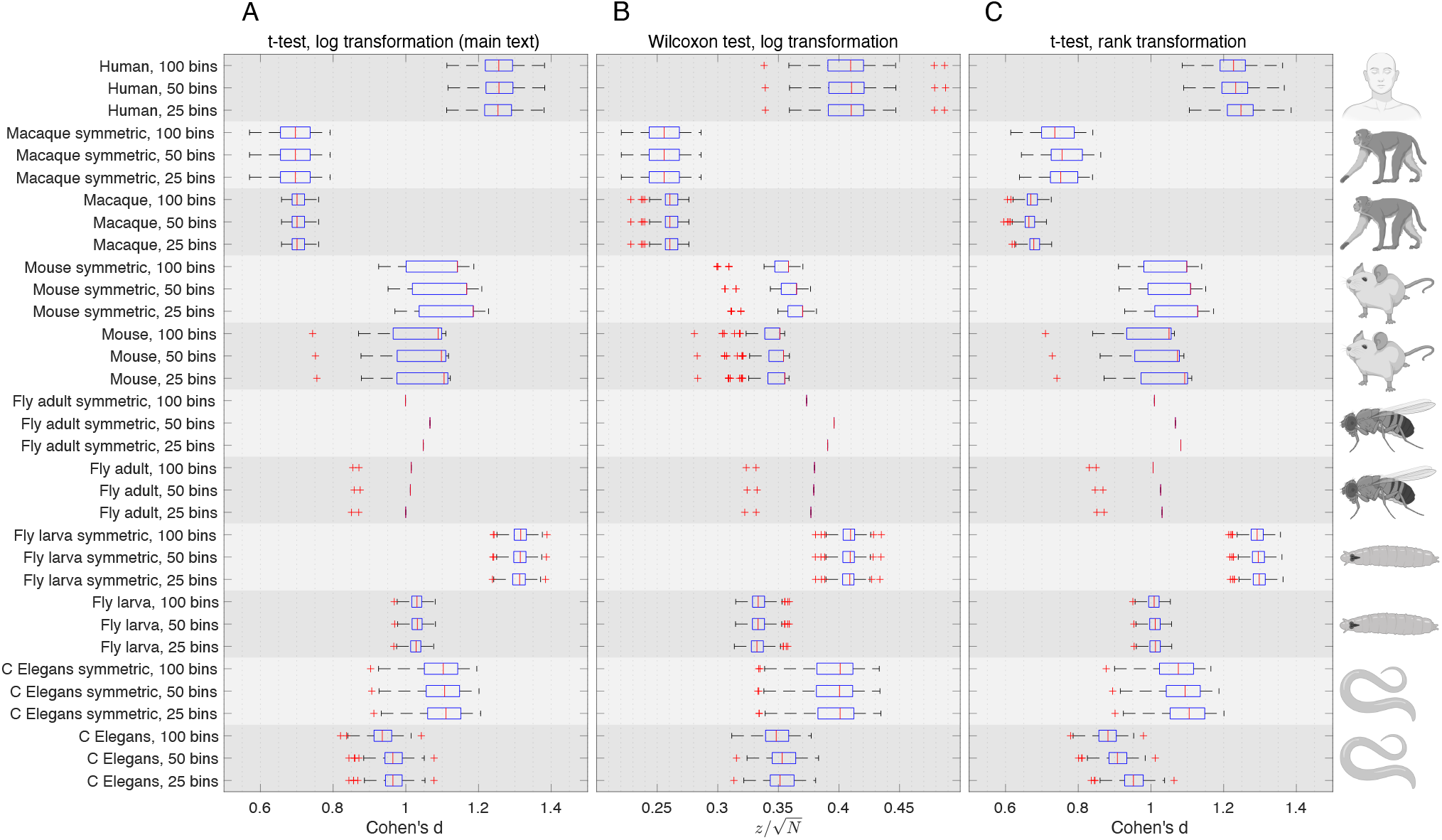
Sensitivity analyses. Horizontal axes show the effect sizes of statistical tests of the difference in within-*vs* between-module RDS. Box plots show the effect size distributions obtained from 100 repetitions of the Louvain algorithm with *γ*=1. The RDS was computed considering 25, 50 (main text parameter), and 100 routing-axis bins, as well as for directed and symmetrised connectomes. A structural connectivity matrix *W* was symmetrised as *W*_sym_ = (*W* + *W* ^*t*^)*/*2. Statistical tests were computed using (A) two-sample *t*-tests and log-transformed communication measures (main text parameters), (B) nonparametric Wilcoxon rank sum test and log-transformed communication measures, and (C) two-sample *t*-tests and ranktransformed communication measures.

